# DSS-induced inflammation in the colon drives a pro-inflammatory signature in the brain that is ameliorated by prophylactic treatment with the S100A9 inhibitor paquinimod

**DOI:** 10.1101/2021.09.22.461244

**Authors:** Sarah Talley, Rasa Valiauga, Lillian Anderson, Abigail R. Cannon, Mashkoor A. Choudhry, Edward M. Campbell

**Author notes:** Corresponding author Edward Campbell.

## Abstract

**Background:** Inflammatory Bowel Disease (IBD) is established to drive pathological sequelae in organ systems outside the intestine, including the central nervous system (CNS). Many patients exhibit cognitive deficits, particularly during disease flare. The connection between colonic inflammation and neuroinflammation remains unclear and characterization of the neuroinflammatory phenotype in the brain during colitis is ill-defined.

**Methods:** Transgenic mice expressing a bioluminescent reporter of active caspase-1 were treated with 2% Dextran Sodium Sulfate (DSS) for 7 days to induce acute colitis, and colonic, systemic and neuroinflammation were assessed. In some experiments, mice were prophylactically treated with paquinimod (ABR-215757) to inhibit S100A9 inflammatory signaling. As a positive control for peripheral-induced neuroinflammation, mice were injected with lipopolysaccharide (LPS). Colonic, systemic and brain inflammatory cytokines and chemokines were measured by cytokine bead array (CBA) and Proteome profiler mouse cytokine array. Bioluminescence was quantified in the brain and caspase activation was confirmed by immunoblot. Immune cell infiltration into the CNS was measured by flow cytometry, while light sheet microscopy was used to monitor changes in resident microglia localization in intact brains during DSS or LPS-induced neuroinflammation. RNA sequencing was performed to identify transcriptomic changes occurring in the CNS of DSS-treated mice. Expression of inflammatory biomarkers were quantified in the brain and serum by qRT-PCR, ELISA and WB.

**Results:** DSS-treated mice exhibited clinical hallmarks of colitis, including weight loss, colonic shortening and inflammation in the colon. We also detected a significant increase in inflammatory cytokines in the serum and brain, as well as caspase and microglia activation in the brain of mice with ongoing colitis. RNA sequencing of brains isolated from DSS-treated mice revealed differential expression of genes involved in the regulation of inflammatory responses. This inflammatory phenotype was similar to the signature detected in LPS-treated mice, albeit less robust and transient, as inflammatory gene expression returned to baseline following cessation of DSS. Pharmacological inhibition of S100A9, one of the transcripts identified by RNA sequencing, attenuated colitis severity and systemic and neuroinflammation.

**Conclusions:** Our findings suggest that local inflammation in the colon drives systemic inflammation and neuroinflammation, and this can be ameliorated by inhibition of the S100 alarmin, S100A9.

## Background

Nearly 1.5 million North Americans and 2.2 million Europeans suffer from inflammatory bowel disease (IBD), and the annual incidence and prevalence continue to increase [1, 2]. IBD is characterized by chronic inflammation in the intestinal tract, consisting of ulcerative colitis (UC) and Crohn’s disease (CD). In UC, inflammation typically involves the rectum and extends proximally, resulting in lesions of variable severity that remain localized to the colon [3]. In CD, inflammation is transmural, producing segmented lesions that can occur in any part of the gastrointestinal tract [3]. Both UC and CD are life-long diseases that cycle between periods of active flare and remission. While pathology primarily affects the intestine, extraintestinal manifestations are prevalent and can involve virtually any organ site [3-5]. In this regard, IBD is increasingly linked to CNS inflammation and behavioral alterations in patients. A relatively large number of IBD patients report depressive symptoms and cognitive dysfunction compared to the general population, ranging from symptoms that seem to be transient and coincide with flare onset to cognitive issues that a recent study reveals as a long-term comorbidity in IBD patients [6-11]. IBD patients are also at increased risk for neurodegenerative diseases, such as Parkinson’s disease and dementia [12-17]. Understanding the mechanistic basis of this connection between colonic inflammatory disease and neurological manifestations will be necessary to develop therapies designed to alleviate or prevent colitis-induced CNS dysfunction.

We have recently developed a transgenic mouse expressing a caspase-1 biosensor that allows the measurement of inflammatory responses in living animals and *ex vivo* [18]. Initial characterization of this mouse demonstrated that the biosensor can effectively identify regions of segmented inflammation in the colon of mice treated with DSS [18]. In this study, we employed our reporter system to monitor inflammation in mice with colitis. Our findings demonstrate that local inflammation in the colon triggers systemic and neuroinflammation. We further characterized this proinflammatory signature in the brains of mice with colitis and found that prophylactic administration of paquinimod can mitigate colonic and CNS inflammation.

## Methods

Mice. IQAD Tg C57Bl/6 mice were bred in-house and maintained in pathogen-free conditions at Loyola University Chicago. Both male and female mice were used for these experiments. All experiments were performed according to protocols approved by the Institutional Animal Care and Use Committee at Loyola University Chicago.

Induction of DSS and clinical measurements of colitis. Mice were given 2% (wt/vol) DSS (40,000 kDa; MP Biomedicals), *ad libitum*, in their drinking water for 7 days. In some experiments, mice were given 2% DSS for 5 days, followed by normal drinking water for 2 days. Control mice received normal water. In some experiments, mice were treated with 10 mg/kg/day paquinimod (MedChem Express), added into the drinking water starting 7 days prior to the addition of DSS and continued during the DSS regimen. Control mice received the same dose of paquinimod for 14 days. In all experiments, water was changed twice a week. Body weight loss was determined and mice were monitored for rectal bleeding, diarrhea and signs of morbidity over the course of disease. On day 7 post-DSS, mice were sacrificed. Tissues were extracted and bioluminescent images were acquired *ex vivo*. Colon length was measured and blood in the stool was assessed using a hemoccult blood test kit (Beckman Coulter). Colon sections were fixed in 10% phosphate buffered formalin. Sections were paraffin embedded, processed and stained with H&E by AML laboratories (Saint Augustine, FL). Blood was collected by submandibular bleeding from the cheek on day 5 or cardiac puncture on day 7. Tissues were harvested and processed as described below.

IVIS imaging. Mice were injected with 150 mg/kg VivoGlo Luciferin and sacrificed 10 minutes later. Tissues were extracted and imaged using the IVIS 100 Imaging System (Xenogen). Prior to imaging, colons were flushed with PBS, followed by 300 ug/mL luciferase substrate. Regions of interests were drawn to gate around the tissue, and bioluminescence (max radiance – p/s/cm2/sr) was quantified within each region using Living Image software.

Organotypic brain slice cultures. Slice cultures were generated as described previously [19], with some modifications. Briefly, brains were removed from control and DSS-treated mice and placed in cold Gey’s buffer solution before slicing 200 μm slices using a vibratome (Leica VT1000S). Slices were transferred to Millicell-CM membrane inserts (0.4 μm pore size, 30 mm diameter, 2-3 slices per insert; Millipore Corp) in six well tissue culture treated plates containing MEM with 25% horse serum and 6.5 mg/mL of glucose. Slices were incubated in 5% CO2 at 37°C for 48 hours before a complete media change to MEM with 20% house serum and 6.5 mg/mL of glucose. Slices were cultured for 7 days with media changes every other day. On day 7, inserts with slices were placed in a new 6 well containing 1 mL VivoGlo luciferin (300 ug/mL) and imaged using the IVIS. Bioluminescence was quantified using Living Image software.

Flow cytometry. Cells were isolated from brain tissue as previously described [20], with some modifications. Briefly, isolated tissues were cut up into fine pieces and were dissociated in buffer containing HBSS (without calcium/magnesium), 5% FBS, 10μM HEPES, 2mg/mL collagenase D (Sigma-Aldrich) and 28U/mL DNaseI (NEB) at 37°C for 45 mins. Every 15 minutes, dissociated tissue was pipetted up and down using sequentially smaller Pasteur pipettes to homogenize tissue debris. Homogenate was filtered through a 70 μm filter and centrifuged 10 mins at 300g. Pellets were resuspended in PBS and stained with an APC-Cy7 LIVE/DEAD^™^ fixable dead cell stain kit (Invitrogen). Cells were washed, resuspended in FACS buffer with Fc block (BD Biosciences) and stained with the following antibodies: AF700 anti-CD3 (BioLegend, clone:17A2), BV711 anti-Ly6C (BioLegend, clone: 1A8), BV421 anti-Ly6G (BioLegend, clone: HK1.4), FITC anti-CX3CR1 (BioLegend, clone: SA011F11), BV510 anti-CD19 (BioLegend, clone: 6D5), PerCPCy5.5 anti-CD45 (BioLegend, clone: 30-F11), and APC anti-Cd11b (BioLegend, clone: M1/70). Samples were measured on an LSRFortessa (BD Biosciences) and analyzed using FlowJo software.

iDISCO+ and light sheet microscopy. Brain tissues were stained and clarified following the iDISCO+ protocol (https://idisco.info/idisco-protocol/). The protocol was executed precisely, with primary and secondary antibody incubations lasting 7 days each following methanol pretreatment. The primary antibody used was anti-Iba1 (Novus Biologicals, cat: NB100-1028) at 2 ug/mL and the secondary antibody was Alexa Fluor 647 AffiniPure Donkey Anti-Goat IgG (Jackson Immunoresearch, cat.: 705-605-147) used at 15 ug/mL. Following clarification, the samples were imaged using LaVision UltraMicroscope II (Northwestern University) equipped with an Andor Neo cMOS camera and a 2XC/0.50 NA objective lens (Olympus) with a 6mm working distance dipping cap. We imaged using the 640-nm laser with a step size of 10um and the continuous scanning light sheet method. Data were analyzed and images generated using Imaris software (Bitplane). Three biological replicates per treatment group were used for this experiment.

Cytokine and chemokine measurements. Tissues were weighed and homogenized using an electric homogenizer. Homogenates were centrifuged at 3900 rpm for 10 minutes to pellet debris, and the supernatant was collected and stored at -80°C until further use. Serum was isolated from whole blood collected from the submandibular vein or by cardiac puncture. Levels of proinflammatory cytokines and chemokines were measured using cytokine bead array (CBA) (BD Biosciences), according to the manufacturer’s protocol. Specifically, samples were incubated with capture and detection beads for a selected panel of cytokines and chemokines – IL-1α, IL-1β, IL-6, TNF, KC, MCP-1, MIP-1α and MIP-1β. Samples were measured on an LSRFortessa (BD Biosciences) and analyzed using FCAP Array software v3.0 (BD Biosciences). The log2FC was calculated by normalizing each sample to the average expression level detected in control samples for each cytokine/chemokine. Binary logarithm (Log base 2) values were calculated and the log2FC values were imported into R to generate heat maps, where 0 = white, red = upregulated, and blue = downregulated expression of the indicated cytokine/chemokine. For the measurement of brain cytokines and chemokines, the Proteome Profiler Mouse Cytokine Array Kit, Panel A (R&D Systems) was used following the manufacturer’s instructions. S100 Calcium Binding Protein A8 and S100 Calcium Binding Protein A9 Heterodimer was measured in tissue homogenates or serum using the Mouse S100A8/S100A9 Heterodimer DuoSet ELISA kit, according to the manufacturer’s protocol (DY8596-05, R&D systems).

Protein isolation and immunoblot. Tissue homogenates were resuspended in 1x lysis buffer (1% NP-40, 100mM Tris, pH 8.0, and 150mM NaCl) containing a protease inhibitor mixture (Roche) and shaken 60 mins at 4°C. Samples were centrifuged and lysate was collected. Protein concentration was quantified using Pierce BCA protein assay kit (Thermo Fisher Scientific). Tissue lysates were mixed with 2x Laemmli sample buffer and incubated at 100°C for 5 min and equal amounts of protein were loaded into a 4-15% polyacrylamide gel for SDS/PAGE and transferred onto a nitrocellulose membrane (Bio-Rad). Membranes were incubated with the following primary antibodies: anti-caspase-1 (p20) (Casper-1, AdipoGen), anti-caspase-3 (#9662, Cell Signaling Technology), anti-caspase-7 (#9492, Cell Signaling Technology), anti-caspase-8 (#8592, Cell Signaling Technology), anti-caspase-8 (sc-81656, Santa Cruz Biotechnology), anti-Lipocalin-2 (AF1857-SP, R&D Systems) and anti-β-Actin (sc-8432, Santa Cruz Biotechnology). Membranes were then probed with anti-mouse, anti-rabbit (Thermo Fisher Scientific) or anti-goat (Promega) secondary antibodies conjugated to horseradish peroxidase, and antibody complex were detected using SuperSignal West Femto Maximum Sensitivity Substrate (Thermo Fisher Scientific).

RNA isolation and qRT-PCR. Tissue homogenates were resuspended in RNA lysis buffer and RNA was isolated using the NucleoSpin RNA Plus RNA isolation kit (Macherey-Nagel). For some experiments, RNA was isolated from brain tissue by TRIzol reagent following the manufacturer’s protocol (Thermo Fisher Scientific). cDNA was synthesized using the GoScript Reverse Transcription System (Promega). Quantitative real-time PCR was conducted using primer sets for the indicated transcripts and iTaq Universal SYBR Green master mix (Bio-Rad).

RNA sequencing and pathway analysis. Mice were treated with 2% DSS for 7 days or 2% DSS for 7 days followed by normal drinking water for 7 days. Control mice received normal water for 7 days. Mice were euthanized on day 7 or 14 and brains were harvested and dissected into three general regions - forebrain, midbrain, and hindbrain. Forebrain sections were washed and total RNA extracted using *mir*Vana miRNA Isolation Kit (Invitrogen), according to manufacturer’s instructions. Total RNA concentration and purity were assessed using a Nanodrop spectrophotometer (ThermoScientific). Isolated total RNA from forebrain samples was submitted to Novogene for library preparation and RNA sequencing analysis. Endotoxin quantification. Endotoxin levels were measured following the manufacturer’s instructions (Pierce LAL Chromogenic Endotoxin Quantification Kit, Thermo Scientific). Specifically, serum isolated from mice was cleared of all RBCs, heat inactivated, diluted 50-fold in endotoxin free water, and plated in duplicate. Absorbance was measured at 405nm.

Statistical analysis. Statistical significance was assessed using the Student’s t test comparing each group to its respective control. *P* < 0.05 was considered significant. Graphs were generated and calculations were performed using GraphPad Prism software (GraphPad Software, Inc.).

## Results

To understand the inflammatory response in mice with ongoing colitis, we employed the DSS model of colitis, which recapitulates human UC in mice. Similar to our previous results [18], caspase biosensor mice given 2% DSS for 7 days exhibited significant weight loss, colonic shortening, increased histopathological hallmarks of UC and blood in the stool, as well as biosensor activation and increased inflammatory cytokine/chemokine levels in the inflamed colon (**Fig. 1**). Female mice developed less severe disease, as measured by reduced weight loss, less colonic shortening, reduced inflammatory cytokine and chemokine levels in the colon, and less female mice had a positive Hemoccult test (**Fig. 1, Supplement 1**).

**Figure 1.**
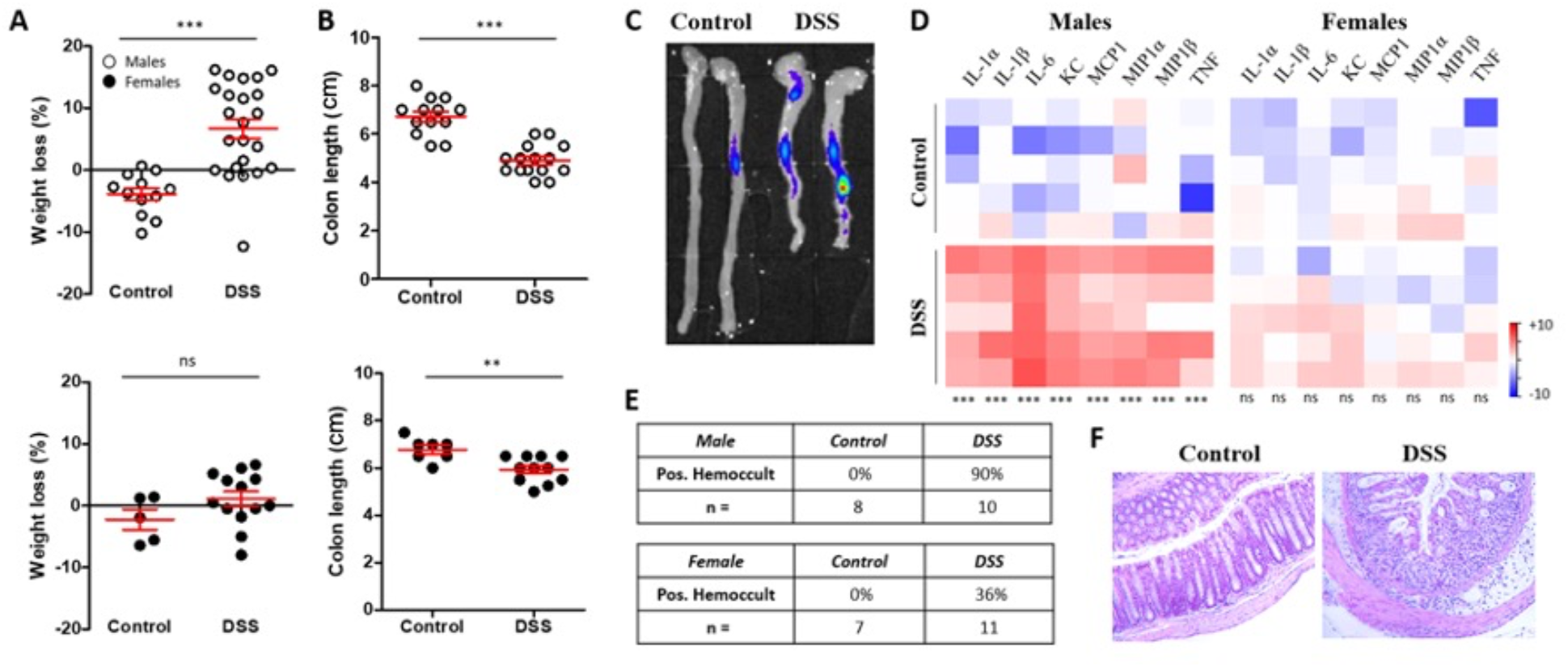
Mice with colitis exhibit inflammation in the colon. Male mice (white circles) and female mice (black circles) were treated with 2% DSS for 7 days. Body weight (A), colon lenghts (B) and caspase biosensor signal (C) were measured. Colon sections isolated from control and DSS mice were homogenized and inflammatory cytokine and chemokine levels were measured by CBA in control or DSS-treated male mice (D). CBA data are expressed as the log2FC in cytokine/chemokine expression, relative to the average cytokine/chemokine levels measured in control mice. Representative expression data from 5 mice in each group is shown (D). Fecal samples were isolated from control and DSS-treated mice and assessed for blood in the stool by Hemoccult (E). Representative colonic H&E sections from one control male and one DSS-treated male mouse (F). Student’s t test comparing DSS to control. *** = p<0.0005, ** = p<0.005.

We next tested whether the induction of colitis could drive inflammasome activation in other tissues, specifically the CNS. We observed a significant increase in biosensor activation in the brains of mice treated with DSS compared to control mice (**Fig. 2A**). We were concerned that a disruption of the blood brain barrier (BBB) could lead to increased luciferase substrate diffusing into the brains of mice treated with DSS, and therefore may explain the apparent increase in biosensor activation observed in the brains of these mice. To address this concern, we generated organotypic slice cultures from brains isolated from DSS-treated and control mice immediately following *ex vivo* IVIS imaging. After 7 days of standard culture conditions, bioluminescence was quantified *in vitro*, and we again observed increased biosensor activation in slice cultures generated from DSS-treated mice, relative to control mice (**Fig. 2B**).

**Figure 2.**
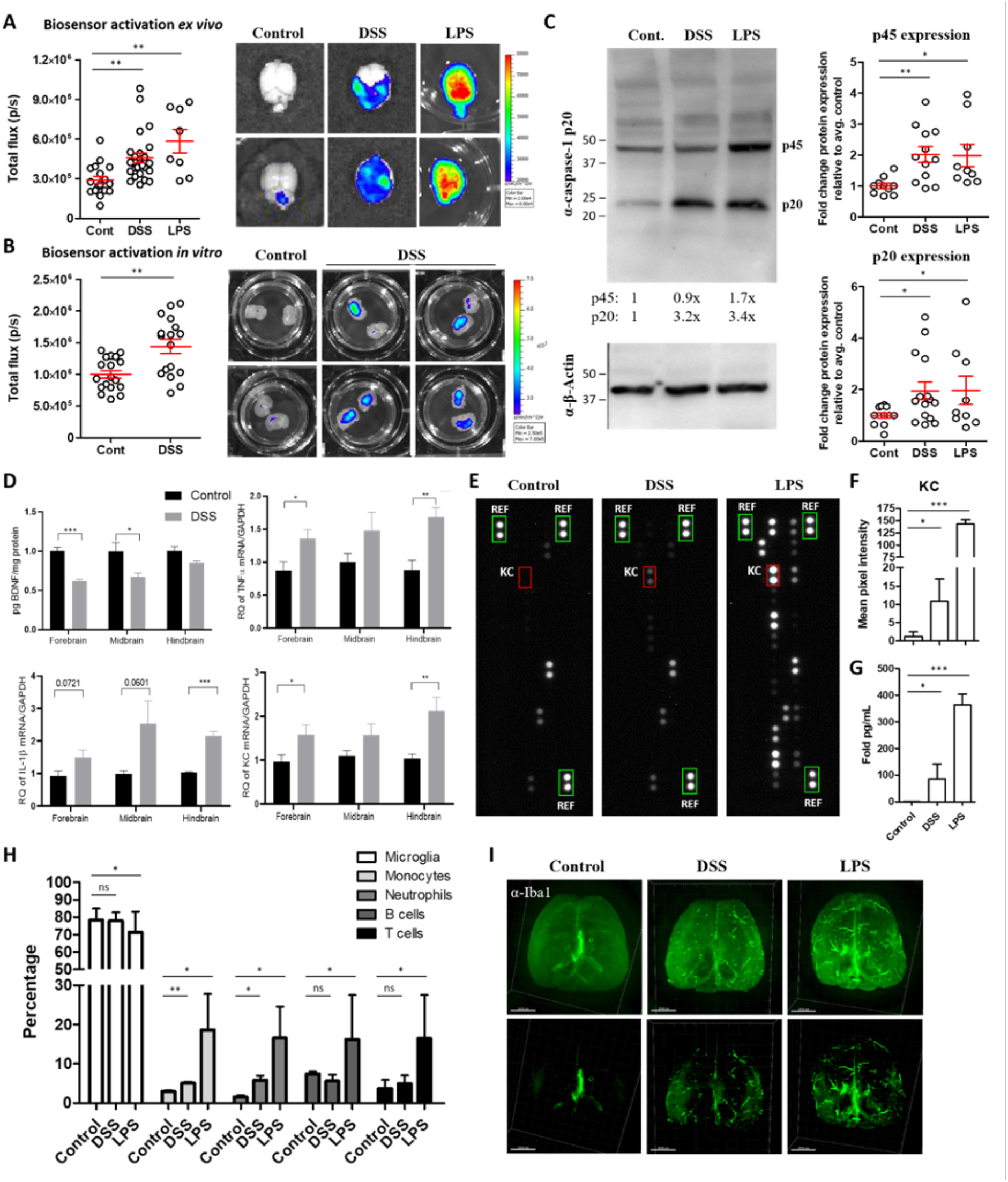
DSS-treated mice exhibit increased neuroinflammation but not immune cell infiltration in the brain. Bioluminescence was quantified from brain tissue harvested from caspase reporter mice either treated with normal drinking water (control) or 2% DSS for 7 days, or injected with LPS (100μg, 24 h) (A). 200μm organotypic slice cultures were generated from control or DSS brains and after one week *in vitro*, luciferase signal was measured using the IVIS (B). Brains were isolated from control or DSS mice, or mice injected i.p with 100μg LPS for 24h. Tissues were homogenized and caspase-1 expression was measured in lysates by western blot (C). Whole brains were sectioned into the indicated regions and BDNF, TNFα, IL-1β and KC levels were measured by ELISA or qRT-PCR, n = 5-8 animals/group (D). Brains were homogenized, lysed and equal amounts of protein lysates were incubated with the mouse cytokine detection antibodies and applied to nitrocellulose membranes pre-coated with 40 different anti-cytokine antibodies in duplicate and protein expression was measured by chemiluminescence (E). Green boxes indicate reference control. Mean pixel intensity of KC (red box) was quantified using ImageJ (F). CBA was performed on brain homogenates and KC expression was quantified (G). Brains isolated from control, DSS and LPS-treated mice were digested and isolated cells were stained for flow cytometric analysis of the indicated immune cell populations, following the gating strategy depicted in Supplement 4 (specifically Live, CD45+, subgated on markers for the indicated immune cell population) (H). Brains were fixed, stained with anti-Iba1 antibody, clarified and imaged using a light sheet microscope (I). Representative images from each group, low min signal panels on the top and high min signal panels on the bottom (I). Student’s t test comparing DSS to control. *** = p<0.0005, ** = p<0.005, * = p<0.05.

Because the bioluminescent reporter can be turned on downstream of either inflammatory and apoptotic caspases [18], we next wanted to determine whether brain tissue isolated from DSS-treated mice exhibited an inflammatory phenotype. Brains were isolated from control or DSS mice 7 days following DSS exposure and caspase-1 expression was measured. We detected an increase in caspase-1 p45 and p20 expression in some, but not all, brains isolated from mice with ongoing colitis (**Fig. 2C**). As a positive control for neuroinflammation induced by a peripheral stimulus, mice were injected intraperitoneally with lipopolysaccharide (LPS) to trigger inflammation in the CNS. We detected significant bioluminescence in brain tissue 24h following LPS injection (**Fig. 2A**). As with the DSS-treated mice, we also measured increased caspase-1 expression in brains isolated from some, but not all, LPS-injected mice (**Fig. 2C**). Notably, in the same DSS and LPS brains where we detected cleaved caspase-1 (p20), we did not detect cleaved caspase-3 (p17/p19), although we did measure some cleaved caspase-8 (**Supplement 2**), which could suggest that other caspases may be contributing to biosensor activation in this system. To further elucidate if inflammatory pathways were upregulated in the CNS in mice with colitis, brain tissue isolated from DSS-treated mice was separated into forebrain, midbrain, and hindbrain regions and expression of inflammatory transcripts was quantified. DSS treatment induced an upregulation of TNF-α, IL-1β and KC (**Fig. 2D**). We also detected reduced expression of brain-derived neurotrophic factor (BDNF), which is well established to be reduced in response to inflammation in a number of pathological conditions [21] (**Fig. 2D**). Next, we measured inflammatory cytokine and chemokine expression using a proteome profiler array to simultaneously measure the relative expression levels of a panel of 40 inflammatory markers in the brains of control, DSS- and LPS-treated mice. Relative to control mice, DSS-treated mice exhibited increased expression of a number of inflammatory cytokines/chemokines, most notably and consistently, KC (**Fig. 2E-F, Supplement 3**). KC upregulation in the brain was confirmed by CBA (**Fig. 2G**).

In order to determine if this increased inflammatory response was the result of increased immune cell infiltration into the brain, immune cell populations in brain tissue from control and DSS-treated mice were assessed. We detected a significant increase in inflammatory monocytes, neutrophils and lymphocytes in the brains of LPS-treated mice, while in DSS-treated mice, we measured minimal increases in monocyte and neutrophil infiltration and no lymphocyte recruitment into the brain (**Fig. 2H, Supplement 4**). To determine if resident cells in the CNS were driving this inflammatory response in the brain in mice with colitis, we isolated brains from control, DSS- or LPS-treated mice, clarified and stained the intact organs with an anti-Iba1 antibody and visualized microglia using light sheet microscopy. We detected increased Iba1 immunoreactivity and a change in signal distribution in DSS brains compared to control, and this signal was even more pronounced in brains isolated from mice injected with LPS (**Fig. 2I**). Collectively, these data suggest that colonic inflammation activates proinflammatory pathways.

We next performed RNA sequencing analysis to better understand the underlying changes in the brains of mice with colitis. Mice were treated with 2% DSS for 5 days and then given normal drinking water for 2 days. On day 7, brain tissues were harvested from DSS and control mice and RNA was prepared for sequencing. Pathway analysis of the differentially expressed genes identified several significantly enriched terms related to inflammation, including genes involved in the IL-17 signaling pathway, regulation of inflammatory responses, and antimicrobial peptides pathway (**Fig. 3A-B**). A number of the genes identified are well established to play a role in neuroinflammation, such as Ptgs2 (also known as COX-2), the receptor for IL-1α/β, IL1R1, and S100 calcium binding proteins S100A8/A9, endogenous alarmins/DAMPs. We focused on Lcn2, S100A8 and S100A9, which were upregulated in 3 of the 4 pathways identified. Lcn2 is an antimicrobial peptide that is secreted by various cell types and has become a biomarker for inflammation. Notably, it is increased in inflamed intestinal tissues of patients with active ulcerative colitis and active Crohn’s disease [22, 23] and our data demonstrate that it is also highly upregulated in the brain of mice with ongoing colitis. Likewise, S100A8 and S100A9, also known as myeloid related protein 8 and 14 (MRP8 and MRP14), respectively, are emerging preclinical biomarkers upregulated in various inflammatory disease models including IBD [24-28]. To this end, we quantified S100A9 and Lcn2 transcript and protein levels in brain tissue isolated from control and DSS-treated mice, using the original experimental paradigm (2% DSS, 7 days). We confirmed upregulation of Lcn2 and S100A9 transcripts and measured increased Lcn2 and S100A8/A9 protein expression in brains isolated from mice with colitis, albeit expression is markedly less than the levels detected in mice injected with LPS (**Fig. 3C-D**). Together, these data reveal transcriptomic changes in the brain consistent with an inflammatory response in the CNS following colitis induction. Importantly, this inflammatory phenotype appears to be transient, as most of the differentially expressed genes detected in the brain on day 7 returned to baseline by day 14, 9 days following DSS cessation (**Fig. 3B**).

**Figure 3.**
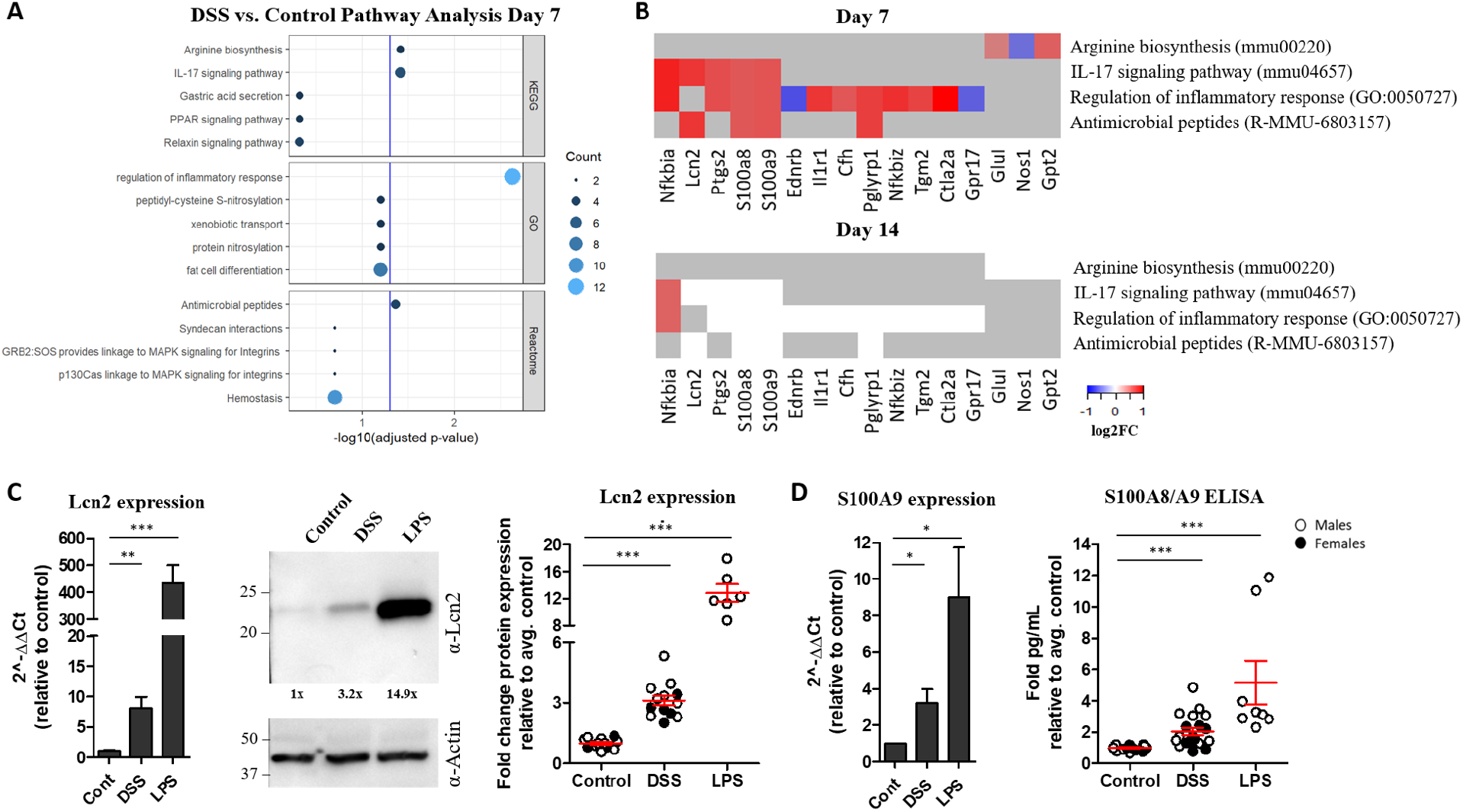
Differential gene expression in brain tissue isolated from mice with colitis reveal an inflammatory signature. Pathway analysis was performed on the differentially expressed genes in the brains of mice treated with DSS compared to control. Significantly enriched terms (adjusted p-value < 0.05) in the KEGG, GO, and Reactome databases are shown (A-B). Genes specifically involved in each term are shown in B. Heatmaps depict the log2FC with up-regulated genes shown in red, down-regulated genes shown in blue, and unchanged genes shown in white (B). Grey represents genes not involved in that pathway (B). Brain tissue was isolated from male and female control mice, mice treated with 2% DSS for 7 days or 100μg LPS for 24h. RNA and protein extracted from homogenized tissues were used for qRT-PCR, ELISA or WB to measure Lcn2 (C), S100A9 and S100A8/A9 (D) expression levels in the brain. For qRT-PCR data, n = 4-13 mice per group. Student’s t test comparing DSS or LPS to control. *** = p<0.0005, ** = p<0.005, * = p<0.05.

In order to determine if there was a systemic inflammatory response in mice with colitis, we measured inflammatory cytokines in the serum. We detected a notable increase in serum IL-6 and KC in mice treated with DSS for 5-7 days (**Fig. 4A-B, Supplement 5**), supporting the idea that acute colitis promotes a systemic inflammatory response [29, 30]. We also measured increased S100A8/A9 in the serum of DSS-treated mice (**Fig. 4C**). However, consistent with previous reports [29], we were unable to detect a significant increase in endotoxin levels in the serum of mice with colitis, despite measuring high levels of serum endotoxin in LPS-treated mice (**Fig. 4D**).

**Figure 4.**
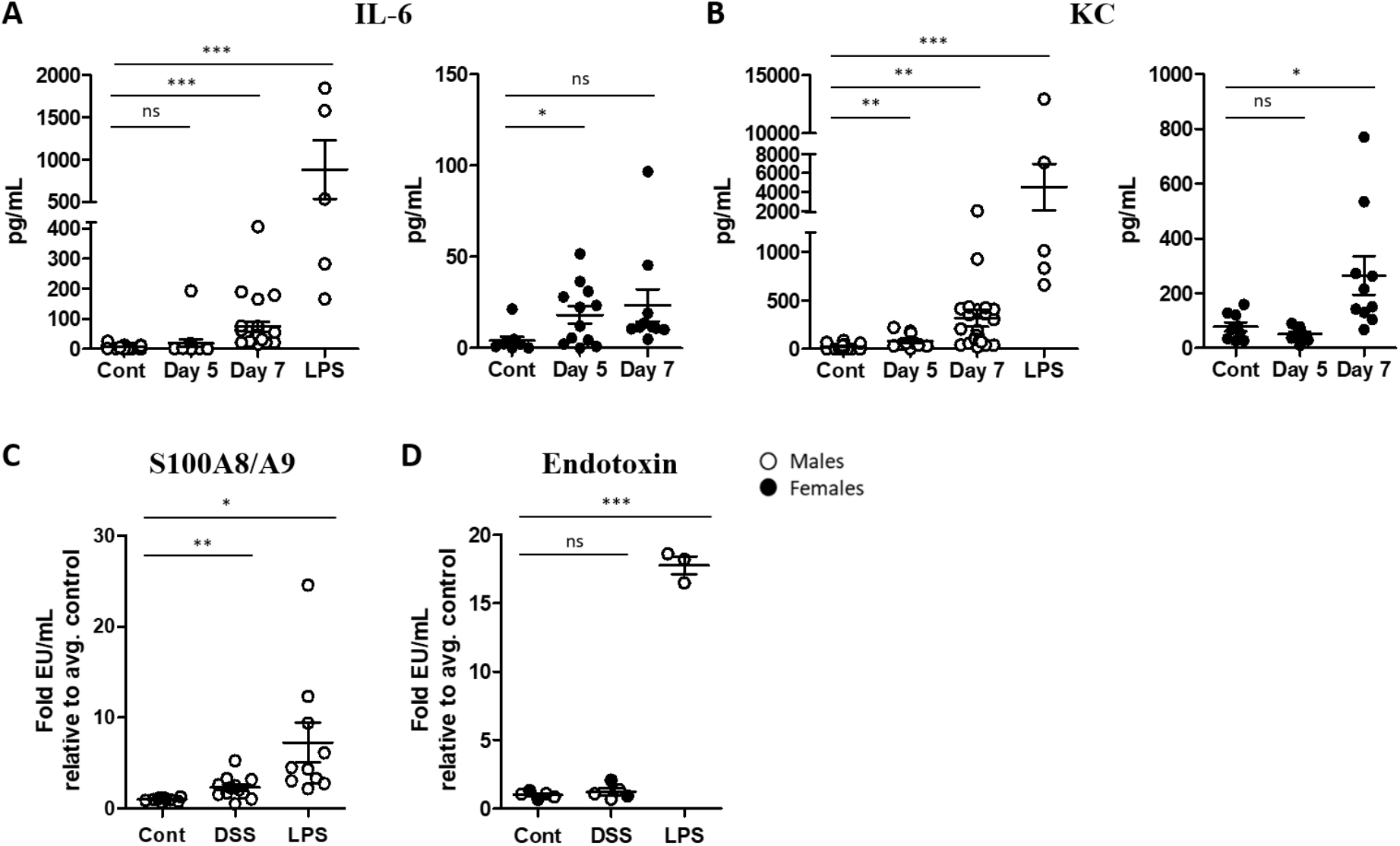
DSS-treated mice exhibit increased inflammatory markers in the serum by day 7 but endotoxin levels remain low. Serum was isolated from male and female control and DSS-treated mice at the indicated time points or mice injected with 100μg LPS for 24h. IL-6 and KC levels were quantified by CBA and S100A8/A9 levels were measured by ELISA (A-C). Serum isolated on day 7 from DSS-treated mice or 24h after injection with 0.5 mg/kg LPS was inactivated, diluted 50-fold in endotoxin-free water and endotoxin levels were quantified (D). Student’s t test comparing DSS to control. *** = p<0.0005, ** = p<0.005, * = p <0.05.

S100A8 and S100A9 are upregulated in numerous models of neuroinflammation, cognitive dysfunction and neurodegeneration [31-34]. Recently, a class of orally-active chemical compounds called quinolone-3-carboxamides (Q compounds) were found to be protective in several inflammatory disease mouse models [32, 35-43] and have been tested in clinical trials [44-46]. One quinolone-3-carboxamide derivative, called paquinimod (ABR-215757), binds S100A9 and inhibits its interaction with TLR4/MD2 and RAGE [47]. To determine if pharmacological inhibition of S100A9 signaling attenuated DSS-induced systemic and neuroinflammation, paquinimod was provided in the drinking water one week prior to and during the onset of colitis. Mice prophylactically treated with paquinimod exhibited detectable but mitigated signs of colitis, as determined by reduced weight loss, colonic shortening and inflammation in the colons compared to DSS-only treated mice, although most mice still had detectable blood in the stool (**Fig. 5A-D**). Consistent with the reduced colonic inflammatory cytokines and chemokines, we also measured reduced levels of S100A8/A9 in the colons of DSS-treated mice (**Fig. 5E**). Paquinimod treatment also prevented the upregulation of Lcn2 and S100A8/A9 in the brain in DSS-treated mice (**Fig. 2F-I**). These data indicate that prophylactic administration of paquinimod ameliorated colonic and neuroinflammation in mice treated with DSS.

**Figure 5.**
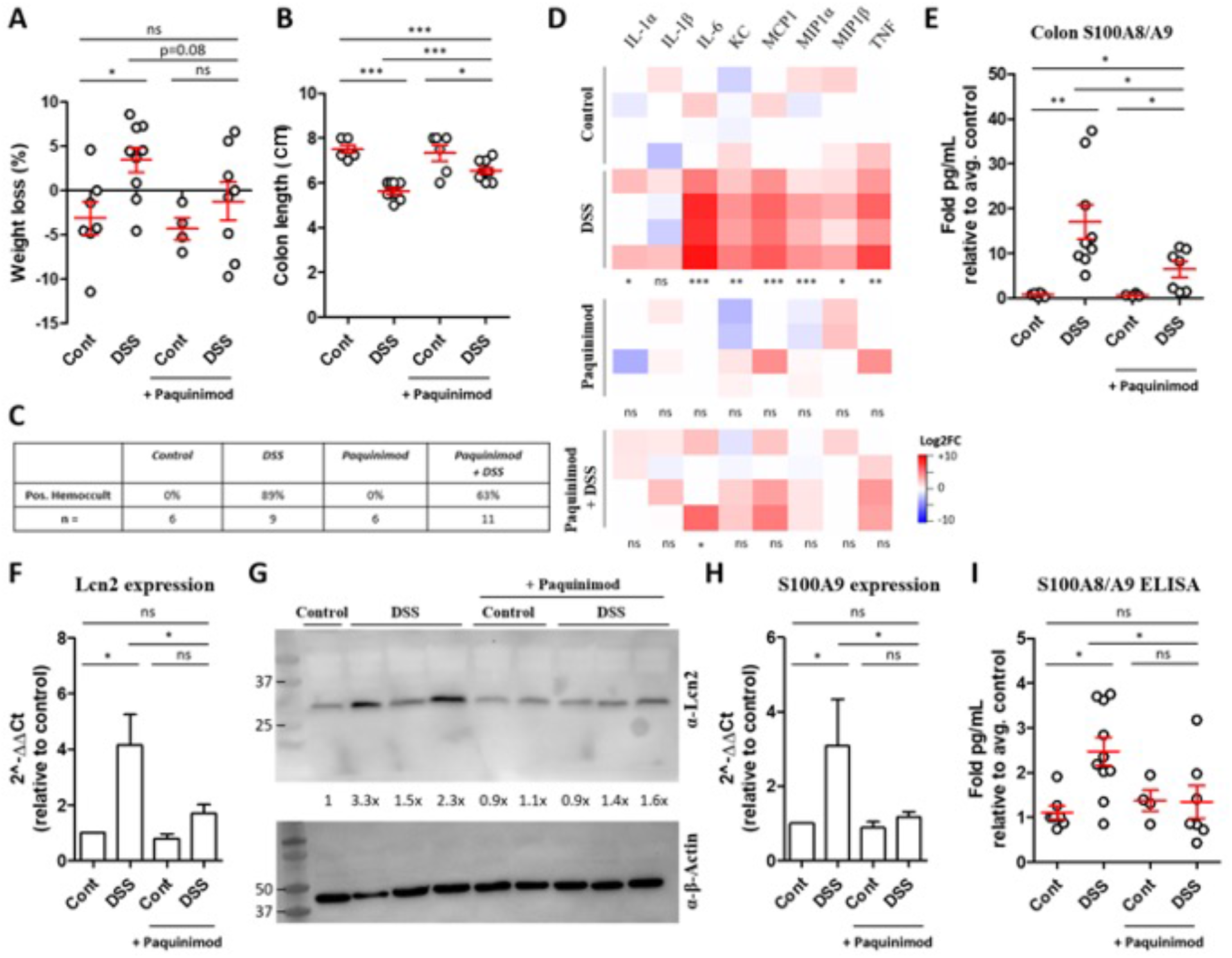
Prophylactic administration of paquinimod reduces inflammation in DSS-treated mice. Male mice were given paquinimod (10mg/kg/day) in the drinking water starting one week prior to DSS. Mice were treated with or without 2% DSS in the absence or presence of paquinimod. 7 days later, weight loss (A), colon lengths (B) and hemoccult (C) were measured. Cytokine/chemokine and S100A8/A9 levels were measured in the colon by CBA (D) and ELISA (E). RNA and protein were isolated from brain tissue homogenates and qRT-PCR was performed to quantify the upregulation of the indicated transcripts (F, H) and proteins (G, I). Student’s t test comparing each treatment group to control (untreated). *** = p<0.0001, ** = p<0.005, * = p<0.05.

## Discussion

Inflammatory manifestations outside of the intestines are common in patients with IBD, and typically coincide with colonic inflammation [5]. To identify extraintestinal sites of inflammation during ongoing colitis, we took advantage of a reporter mouse that becomes bioluminescent at sites of inflammasome/caspase activation. As expected, we detected colonic inflammation and clinical signs of colitis in mice treated with DSS for 7 days (**Fig. 1**). We also detected significant biosensor activation in the brains of mice with colitis (**Fig. 2**). It is important to note that while we have demonstrated that this reporter becomes activated in sites of inflammation, it may also be activated by other inflammatory and apoptotic caspases in addition to caspase-1 [18, 48]. Consistent with increased biosensor activation, we detected increased expression of caspase-1 in brains isolated from DSS and LPS-treated mice (**Fig. 2C**). The increased expression of p45 is likely indicative of increased upregulation of procaspase-1, triggered by the activation of inflammatory signaling cascades, while the increased expression of p20 is a downstream indicator of inflammasome activation. The presence of cleaved caspase-1 together with biosensor activation in the brains of mice treated with DSS is a strong indicator that an inflammasome(s) is active in this tissue during colitis. Identifying the inflammasome-forming receptor(s) that is activated in response to DSS treatment and the cell type or brain region where the inflammasome is activated requires further investigation.

We also detected cleaved caspase-8 in the brains of some DSS- and LPS-treated mice (**Supplement 2**). Caspase-8 activation in microglia has been reported previously *in vitro* in response to challenge with LPS and other inflammatory stimuli [49], as well as in microglia in brain tissue from patients with multiple sclerosis (MS) [50]. In addition to its canonical role as a driver of apoptosis, caspase-8 has recently emerged as a critical player in inflammatory responses. Like caspase-1, active caspase-8 cleaves inflammatory cytokines IL-1β, IL-18, and can be recruited into inflammasome complexes, particularly in cells where caspase-1 is absent or deficient, driving cell death [51-57]. Microglia specifically have been shown to process IL-1β by a NLRP3-ASC-caspase-8-dependent mechanism [50]. While we were unable to detect cleavage of caspase-3 in brains from either group at this time point, we cannot rule out the possibility that this caspase is activated at low levels below the limit of detection in our assay or that caspase-3 may ultimately become activated at later time points. In support of this, one group found increased caspase-3 in the hippocampus of mice treated with 5% DSS for 7 days [29]. Further experiments should be performed to delineate the temporal kinetics of caspase activation in the CNS in mice with colitis.

To confirm this inflammatory phenotype, we measured the upregulation of specific inflammatory transcripts by qRT-PCR and found increased IL-1β, KC and TNF-α particularly in the hindbrain region (**Fig. 2D**), which is the region of the brain that is reproducibly bioluminescent in DSS-treated mice. Proteome mouse cytokine array and CBA also revealed increased expression of a number of inflammatory cytokines and chemokines, most notably KC (**Fig. 2E-G**), which was also significantly elevated in the colon (**Fig. 1D**) and serum (**Fig. 4B**). We next took an unbiased approach to identify differentially expressed genes in the brains of mice with colitis. Pathway analysis of the RNA sequencing data revealed that the most significantly upregulated pathway was the regulation of inflammatory response pathway, followed by IL-17 signaling, arginine biosynthesis and antimicrobial peptides pathways (**Fig. 3**). Inflammatory and antimicrobial pathways are intimately linked, so it is not surprising that both of these pathways were upregulated in our model. We confirmed upregulation of three proteins identified in three of the pathways, S100A8, S100A9 and Lcn2, all of which are established biomarkers for inflammation. Other groups have previously demonstrated that S100A9 [58, 59] and Lcn2 [22, 60, 61] are upregulated in the colon of mice with colitis. However, the finding that they are also increased in the brain in mice with colitis is novel.

Inflammatory responses can occur in diverse cell types in the CNS, though neuroinflammation has mainly been attributed to microglia. LPS-induced microglia activation in the CNS is well supported (reviewed in [62]) but whether colitis drives microglia activation in the rodent brain remains debated [29, 63-65]. Our data revealed that brains isolated from DSS-treated mice show an increase in Iba1 immunoreactivity and signal distribution (**Fig. 2I**). Since Iba1 is a marker of both resting and activated microglia, the increased signal most likely represents a change in microglia morphology and localization rather than an increase in cell number. The Iba1 signal was even more evident in brains isolated from mice 24h after LPS injection and in some cases, appeared to localize around the vasculature (**Fig. 2I**). While light sheet microscopy is not the best system for quantification, the increased Iba1 signal in both DSS and LPS brains is visually apparent and consistent with increased microglial reactivity [66]. Neuroinflammation can also result from disruption of the BBB and infiltration of immune cells into the nervous system, which drive pathological sequelae in a number of disease models. There is some evidence for BBB permeability in trinitrobenzene sulphonic acid (TNBS)-treated rats/rabbits [67, 68] and mild BBB permeability in DSS-treated mice [29, 69]. We therefore wanted to investigate whether peripheral immune cells infiltrated the CNS in mice with colitis. We measured a modest but significant increase in inflammatory monocytes and neutrophils in the brains from mice treated with DSS (**Fig. 2H**). These data suggest that in addition to resident glia cells, innate immune cells may be contributing to neuroinflammation in mice with colitis.

It is worth noting that although our data demonstrate that proinflammatory pathways are upregulated in the CNS during colitis, this proinflammatory signature appears to be transient and moderate. Our RNA sequencing data that identified upregulation of inflammatory pathways 7 days after DSS exposure also revealed that these genes returned to baseline by 9 days following DSS cessation, indicating that these inflammatory responses resolve upon the resolution of colitis. We also included mice injected systemically with LPS as a positive control for neuroinflammation induced by a peripheral stimulus in our experiments to provide scale. While the proinflammatory phenotype is significantly and reproducibly detected by multiple measurements of inflammation in the brains of DSS-treated mice, it is a mild to moderate inflammatory response relative to that detected in the brains of LPS-treated mice (**Figs. 2-4, Supplement 3-4**).

Mechanistically, how colonic inflammation can trigger neuroinflammation has yet to be elucidated. Colonic inflammation likely drives neuroinflammation either by a directed gut-brain signaling axis, potentially mediated by the gut microbiota or afferent neurons, or inflammation in the colon could trigger systemic inflammation that subsequently activates inflammatory pathways in the CNS. These two pathways may not be mutually exclusive. Our data demonstrate a significant increase in the inflammatory chemokine, KC, as well as inflammatory biomarkers, S100A8 and S100A9, in the colons, serum and brains of mice with colitis (**Figs. 1-5**). As mentioned above, colitis-induced extraintestinal inflammation is well documented in mice and in humans. This, taken together with the increase in inflammatory markers in both the serum and the brain argue in favor of a model whereby colitis drives a global inflammatory response that culminates in inflammation in the CNS. More mechanistic insight into how gut driven inflammation contributes to neuroinflammation is needed.

There has been recent interest in S100A8 and S100A9 because they are found at such high concentrations at sites of inflammation. S100A8 and S100A9 make up nearly half of the total cytoplasmic proteins in neutrophils and are significantly upregulated by macrophages in response to inflammatory stimuli. Once released into the extracellular environment, they activate immune cells and vascular endothelial cells, perpetuating inflammation in tissues. Intriguingly, upregulation of these proteins does not appear to be specific for a particular inflammatory disease, as S100A8 and S100A9 have been implicated in a number of seemingly unrelated inflammatory conditions, including rheumatoid arthritis [70-74], CD [27, 28], systemic lupus erythematosus [75-78], cardiovascular diseases [79-81], psoriasis [82, 83], MS [84, 85], and others. More recently, S100A8/A9 was found to be upregulated in patients with COVID-19 and expression correlated with disease severity [86, 87].

Paquinimod is an orally-active immunomodulatory quinoline-3-carboxamide derivative that binds to S100A9 (and S100A12 [88]), and prevents its interaction with TLR4 and RAGE receptors to prevent inflammatory signaling [47]. Paquinimod and related Q-compounds have been tested in clinical trials for a number of human autoimmune and inflammatory diseases, including CD [44-46, 89], and paquinimod specifically has been shown to be protective in mouse models of infection and autoimmune-induced neuroinflammation [32, 35] as well as depression [31]. Because we detected S100A8/A9 upregulation in the serum and brains in mice with colitis (**Fig. 3-4**), we hypothesized that inhibition of S100A9-signaling would reduce the neuroinflammatory signature in mice with colitis. To determine the effect of S100A9 inhibition on colitis-induced inflammation, mice were prophylactically treated with paquinimod 1 week prior to and during DSS treatment. Paquinimod-treated mice exhibited reduced inflammatory cytokines and chemokines in the colon and reduced inflammatory biomarkers in the brain (**Fig. 5**). Paquinimod and DSS-treated mice also exhibited detectable but improved clinical signs of colitis (**Fig. 5**). These data are supported by the previous finding that treating mice with a neutralizing anti-S100A9 antibody prevented DSS-induced colitis and colitis-associated cancer in mice [58].

The consequence of colitis-induced neuroinflammation on behavior and cognition was not addressed in this work. Previous studies have demonstrated memory impairment, anxiety-like behavior, and altered responses to stress in mouse models of DSS-induced colitis [30, 90-93], which return to normal as acute colitis resolves [92, 93]. Systemic LPS administration has been shown to drive hypothalamic, hippocampal and cortical proinflammatory cytokine expression, and elevation of proinflammatory cytokines in the brain have been linked to memory impairment and neurodegeneration [94-99]. The cognitive deficits and increased stress/anxiety measured in DSS-treated mice and in humans may be, in part, due to this inflammatory milieu in the CNS that persists during colonic inflammation. Stress-induced upregulation of S100A8 and S100A9 has also been reported previously in mouse hippocampus [100], though it is unclear if S100A8/A9 expression in the CNS is beneficial or detrimental to stress and anxiety. In two separate mouse models, one group found that that upregulation of S100A8/A9 may be protective against anxiety-like behavior following sepsis [101], while another group found that pharmacological inhibition of S100A9 with paquinimod effectively ameliorated depressive behaviors in a model of chronic unpredictable mild stress (CUMS) [31]. It will therefore be important to assess the effect of S100A9 inhibition on DSS-induced stress and anxiety.

One caveat with this study is that broadly targeting inflammatory pathways will not only prevent neuroinflammation but also colonic inflammation, the driver of neuroinflammation in our model. In other words, although we detected reduced systemic and neuroinflammation in paquinimod-treated mice, this could be because most mice never developed significant colitis. It will be important to determine if therapeutic administration of paquinimod mitigates systemic and neuroinflammation in mice that have ongoing colitis. However, we were unable to test this, because mice with colitis often stop drinking their water, meaning that drugs given *ad libitium* may not be taken at the proper dose, if at all. An oral gavage route of administration could be tested in the future but was avoided in this study since this approach can induce stress and weight loss, which would likely affect measurements of neuroinflammation and colitis clinical scores. In humans, however, drugs that target S100A9 could be given after colitis onset as soon as patients become symptomatic. It would be interesting to test such therapeutics to determine if they alleviate colitis and other inflammatory sequelae in patients with IBD.

## Conclusions

In conclusion, this study found that colitis drives mild to moderate inflammation in the brain and increased inflammatory cytokines in the serum, and that colonic and neuroinflammation were mitigated by prophylactic treatment with the S100A9 inhibitor paquinimod.

## Figures

**Supplement 1.**
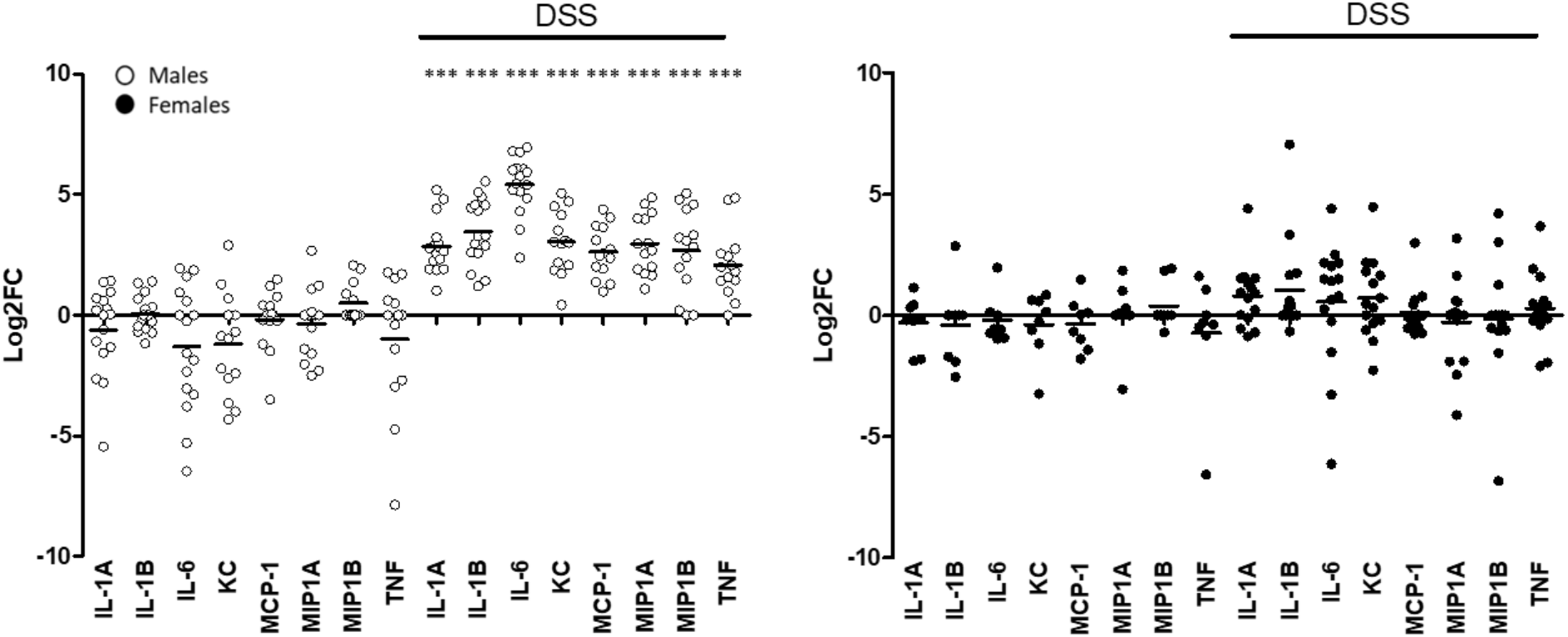
Inflammatory cytokines/chemokines detected in male mice treated with DSS. Mice were treated with 2% DSS for 7 days. Colons were isolated, homogenized and levels of the indicated inflammatory cytokines/chemokines were measured by CBA. The full CBA dataset for male (white) and female (black) is presented, where each individual dot depicts the log2FC of cytokine/chemokine expression in one mouse. Student’s t test comparing DSS to control for each cytokine/chemokine. *** = p<0.0005.

**Supplement 2.**
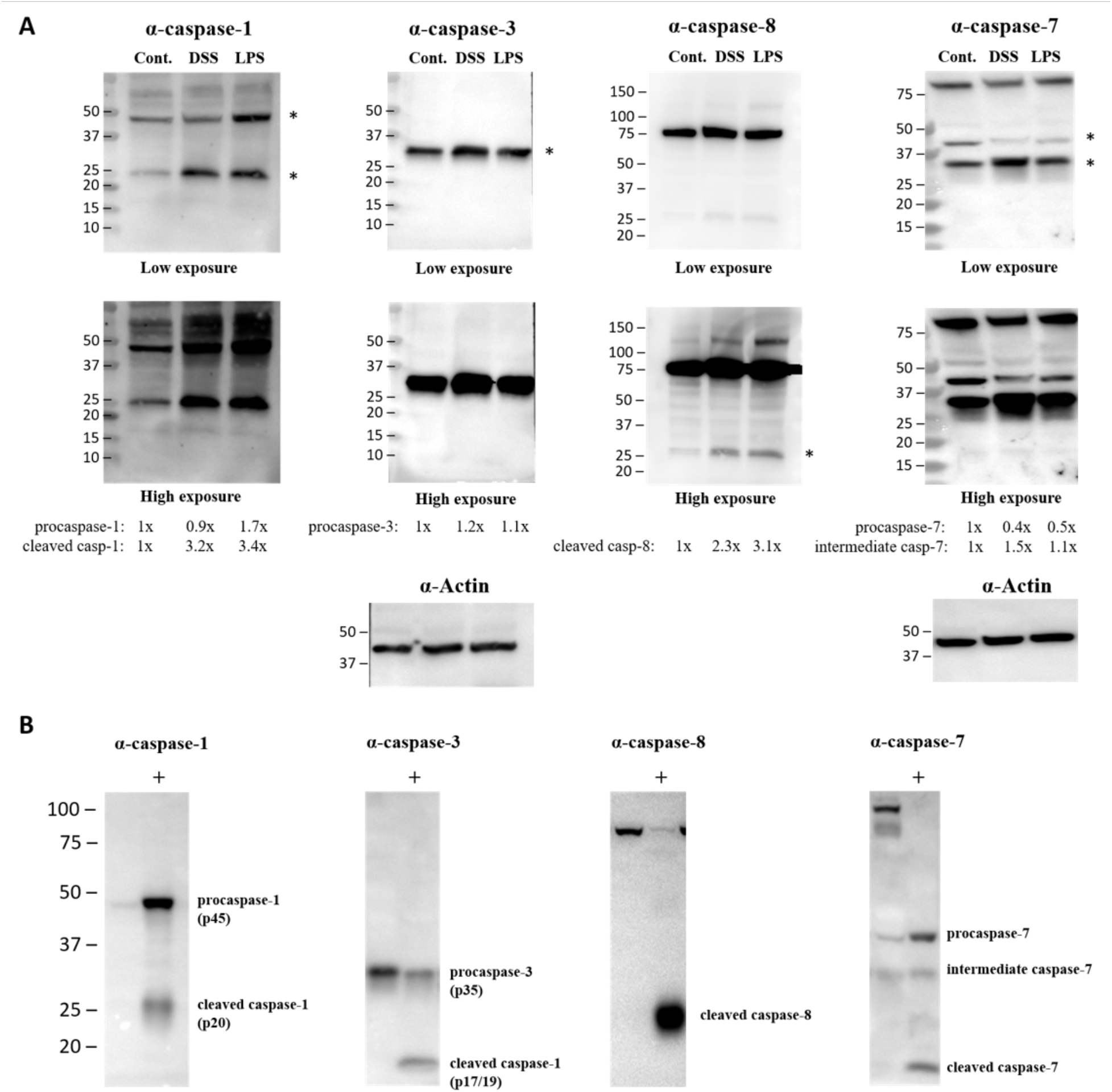
Caspase expression in the brain. Lysates from digested brains isolated from control, DSS or LPS-treated mice were stained with antibodies against caspase-1 p20 (shown in **Fig. 2C**), caspase-3, caspase-8, caspase-7 or actin (A). Bands labelled with an asterisk were quantified (A). Mouse splenocytes treated with 10μM staurosporine *in vitro* were used as a positive control (+ lane) to measure cleaved caspases (B).

**Supplement 3.**
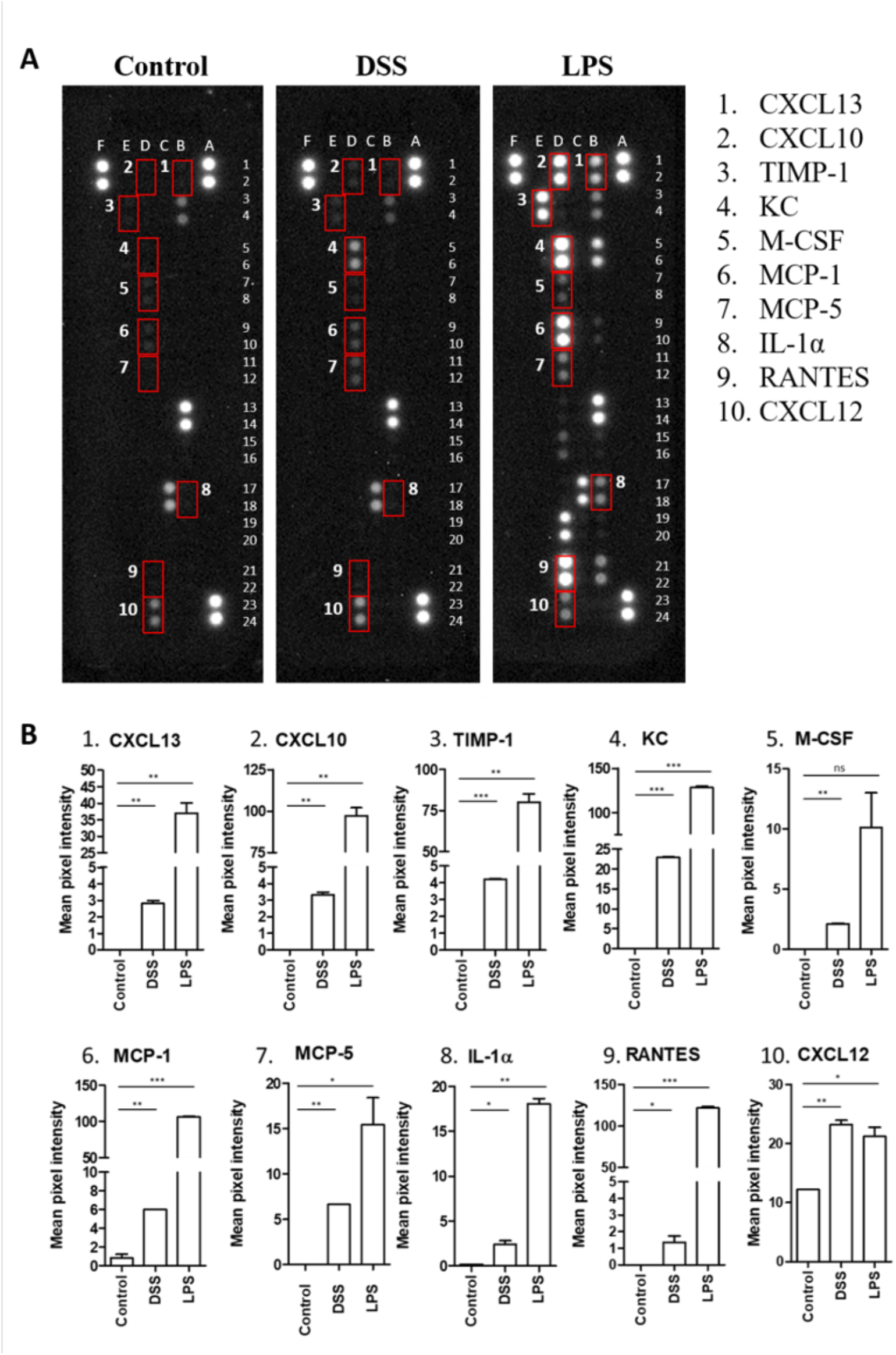
Cytokine/chemokine levels in DSS and LPS-treated brains. Male mice were treated with 2% DSS for 7 days or injected with 100μg LPS for 24h. Brain tissue was collected from control, DSS-treated and LPS-injected mice and ∼10,000 μg whole brain homogenates were incubated with the capture antibody mixture and applied to membranes spotted with detection antibodies for 40 inflammatory cytokines / chemokines (in duplicate). The representative blots (1 control, 1 DSS and 1 LPS) shown in Figure 2 are overexposed to better visualize qualitative changes in expression, and the red boxes indicate the 10 cytokines / chemokines that were significantly upregulated in the brain isolated from the DSS-treated animal (A). Mean pixel intensity was quantified for each spot using ImageJ, and expressed relative to control (B). Student’s t test comparing DSS or LPS to control. *** = p<0.0005, ** = p<0.005, * = p<0.05.

**Supplement 4.**
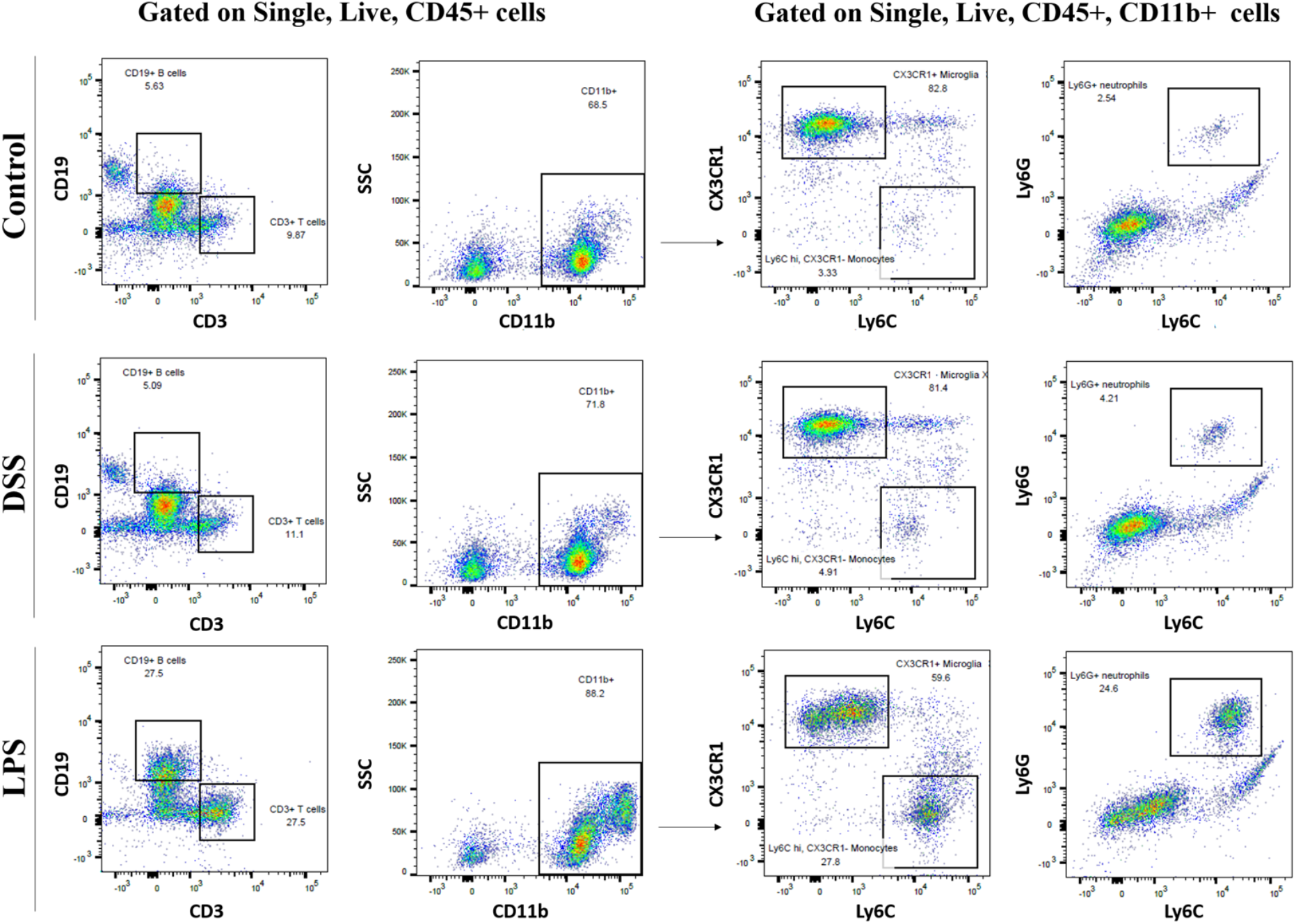
Flow cytometric analysis of immune cell populations in the brain in mice with colitis. Male mice were treated with 2% DSS for 7 days or injected with 100μg LPS for 24h. Brain tissue was collected from control, DSS-treated and LPS-injected mice. Brain tissue was digested and isolated cells were stained for the indicated markers. Single, live, CD45+ cells were gated and CD11b, CD3 and CD19 expression was assessed. Within the CD11b-positive population, CX3CR1, Ly6C and Ly6G expression was assessed. Representative flow plots are shown from a control, DSS and LPS brain. Data are representative of 3 independent experiments. Results were analyzed using FlowJo software.

**Figure S5.**
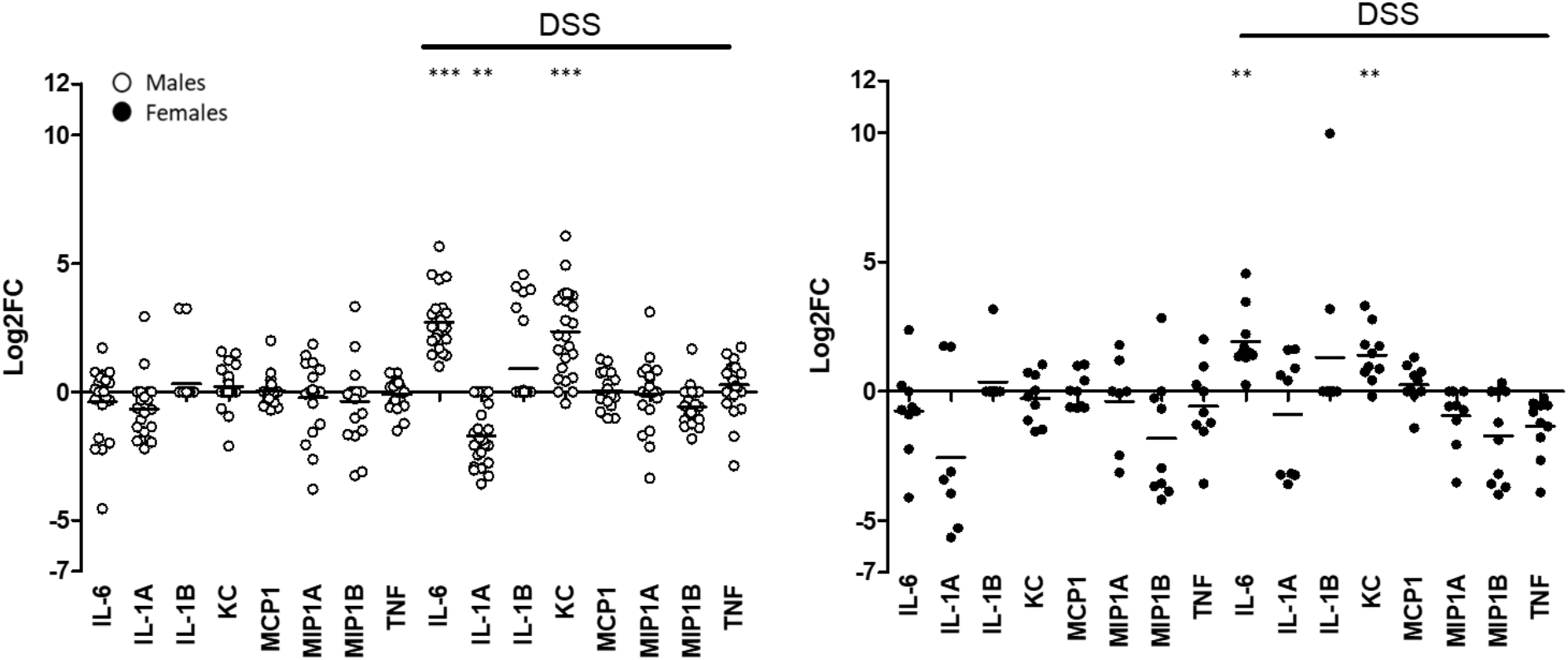
Supplement 5 – Changes in inflammatory cytokines in serum from DSS-treated males and females. Blood was collected and CBA was performed on the serum from control and DSS-treated mice. Dataset depicts the log2FC in cytokine/chemokine expression from male (white) and female (black) mice. Student’s t test comparing DSS to control for each cytokine/chemokine indicated on the x-axis. *** = p<0.0001, ** = p<0.005

## List of abbreviations

IBD: Intestinal bowel disease
CNS: Central nervous system
DSS: Dextran sodium sulfate
LPS: Lipopolysaccharide
CBA: Cytokine bead array
UC: Ulcerative colitis
CD: Crohn’s disease
BBB: Blood brain barrier
IVIS: In vivo imaging system
LPS: Lipopolysaccharide
S100A8/A9: S100 calcium binding proteins A8/A9
Lcn2: Lipocalin-2
RAGE: Receptor for advanced glycation end products
TLR4: Toll-like receptor 4

## Declarations

### Author information

Department of Microbiology and Immunology, Loyola University, Chicago Sarah Talley, Edward M. Campbell

Department of Neuroscience, Loyola University, Chicago Rasa Valiauga, Edward M. Campbell

Stritch School of Medicine, Loyola University, Chicago

Rasa Valiauga

Burn and Shock Trauma Research Institute, Alcohol Research Program, Stritch School of Medicine, Loyola University Chicago Health Science Division Lilian Anderson, Abigail R. Cannon, Mashkoor A. Choudhry

## Ethics approval and consent to participate

All animal work was performed in compliance with the Institutional Animal Care and Use Committee (IACUC) at Loyola University, Chicago (protocol approval numbers 2021003, 2017032, 2019009, 2020023).

## Authors’ contributions

ST, EC and MC conceptualized the study and designed experiments. ST, RV, LA, AC carried out experiments. ST analyzed the data and wrote the manuscript, with guidance from EC and MC. All authors read and approved the final manuscript.

## Acknowledgements

We thank Jessica Mattick (Loyola University Chicago) for assisting with the RNA sequencing and pathway analysis. We thank Costadina Arvanitis (Northwestern University) for assisting with Light Sheet Microscopy imaging.

